# Loss of *Prm1* leads to defective chromatin protamination, impaired PRM2 processing, reduced sperm motility and subfertility in male mice

**DOI:** 10.1101/2021.10.29.466452

**Authors:** Gina Esther Merges, Julia Meier, Simon Schneider, Alexander Kruse, Andreas Christian Fröbius, Klaus Steger, Lena Arévalo, Hubert Schorle

## Abstract

One of the key events during spermiogenesis is the hypercondensation of chromatin by substitution of the majority of histones by protamines. In humans and mice, protamine 1 (*PRM1/Prm1*) and protamine 2 (*PRM2/Prm2*), are expressed in a species-specific ratio. Using CRISPR-Cas9-mediated gene editing we generated *Prm1*-deficient mice and demonstrate, that *Prm1+/-* mice are subfertile while *Prm1-/-* are infertile. *Prm1*-deficiency was associated with higher levels of 8-OHdG, an indicator for reactive oxygen mediated DNA-damage. While *Prm1+/-* males displayed moderate increased levels of 8-OHdG virtually all sperm of *Prm1-/-* males displayed ROS mediated DNA damage. Consequently, DNA integrity was slightly hampered in *Prm1+/-*, while DNA was completely fragmented in *Prm1-/-* animals. Interestingly CMA3 staining which indicates protamine-free DNA revealed, that *Prm1+/-* sperm displayed high levels (93%), compared to *Prm2+/-* (29%) and WT (2%) sperm. This is not due to increased histone retention as demonstrated by mass spectrometry (MassSpec) of nuclear proteins in *Prm1+/-* sperm. Further analysis of the MassSpec data from sperm nuclear proteome revealed, that only one protein (RPL31) is significantly higher abundant in *Prm1+/-* compared to WT sperm. Comparison of the proteome from *Prm1-/-* and *Prm2-/-* to WT suggested, that there are a small number of proteins which differ in abundance. However, their function was not linked mechanistically to primary defects seen in *Prm1-/-* mice and rather represent a general stress response. Interestingly, using acid urea gels we found that sperm from *Prm1+/-* and *Prm1-/-* mice contain a high level of unprocessed, full-length PRM2. *Prm2* is transcribed as a precursor protein which, upon binding to DNA is successively processed. Further, the overall ratio of PRM1:PRM2 is skewed from 1:2 in WT to 1:5 in *Prm1+/-* animals. Our results reveal that *Prm1* is required for proper processing of PRM2 to produce the mature PRM2 which, together with *Prm1* is able to hypercondense DNA. Hence, the species specific PRM1:PRM2 ratio has to be precisely controlled in order to retain full fertility.

## Introduction

During spermatogenesis in the seminiferous epithelium of the testis diploid spermatogonia differentiate into haploid spermatids. One of the most remarkable changes during spermiogenesis is complete reorganization of chromatin compaction [1], where histones are nearly completely substituted by protamines. These are highly basic, arginine rich proteins [2] which, upon binding to DNA hypercondense chromatin, leading to transcriptional silencing and protection of the paternal genome [3]. While in most mammals, DNA compaction in sperm is accomplished by incorporation of protamine 1 (PRM1) alone, primates and most rodents express two protamines, PRM1 and protamine 2 (PRM2) [4, 5]. In mice and men, *Prm1* and *Prm2* are encoded in a tightly regulated gene cluster on chromosome 16 [6, 7]. While PRM1 is expressed as mature protein, PRM2 is expressed as precursor protein (pre- PRM2), consisting of a C-terminal mature PRM2 (mPRM2) domain and a N-terminal cleaved PRM2 (cPRM2) domain, which is sequentially cleaved off upon binding to DNA [5, 8, 9]. Of note, *mPrm2* is proposed to originate from a gene duplication of *Prm1* [10]. In an evolutionary context *Prm1* and *cPrm2* were shown to be conserved, suggesting important roles in fertility [11, 12]. *PRM1*/PRM1 and *PRM2*/PRM2 are detected in a species-specific ratio (humans 1:1 ([13]), mice 1:2 [14]). In humans, alterations of the protamine ratio (PRM1:PRM2) have been associated with male sub- and infertility [15-27]. Mice chimeric for a deletion of one allele of *Prm1* or *Prm2* [28-30] were infertile and did not allow for the establishment of mouse lines and detailed analysis of *Prm*-deficiency. Further, heterozygous *Prm1*-deficient mice generated with CRISPR-Cas9 have been reported to be infertile [31]. Hence, a detailed phenotypical analysis of *Prm1*-deficient mice was not possible so far.

Schneider *et al*. reported the establishment of *Prm2*-deficient mouse lines using CRISPR-Cas9-mediated gene editing in zygotes [32]. Here, *Prm2+/-* male mice remained fertile while *Prm2-/-* were infertile. While *Prm2+/-* sperm showed no pathomorphological effects, *Prm2-/-* sperm presented with fragmented DNA, disrupted sperm membranes and complete immotility. These defects were shown to accumulate during epididymal transit. It was demonstrated that the *Prm2-/-* mice display a deregulation of proteins leading to an accumulation of reactive oxygen species (ROS) explaining the phenotype observed [33]. Using CRISPR-Cas9-mediated gene-editing in zygotes, we generated mice deficient for *Prm1*. Male mice heterozygous for the mutation (*Prm1+/-*) are subfertile, while *Prm1*-deficient (*Prm1-/-*) males are sterile. Molecular analyses revealed that loss of one allele of *Prm1* leads to a moderate fragmentation of DNA, while in *Prm1-/-* mice complete DNA fragmentation can be observed. Sperm of *Prm1+/-* mice display reduced motility as well as enhanced 8-OHdG levels indicative of upregulated ROS levels. Most importantly, analyses of sperm nuclear proteins revealed that the processing of PRM2 to its mPRM2 form seems disturbed in *Prm1+/-* animals already. Further, the species-specific protamine ratio is shifted in *Prm1+/-* mice. These data strongly suggest that the species-specific level of PRM1 is required for proper sperm function.

## Material and Methods

### Ethics statement

All animal experiments were conducted according to the German law of animal protection and in agreement with the approval of the local institutional animal care committees (Landesamt für Natur, Umwelt und Verbraucherschutz, North Rhine-Westphalia, approval ID: AZ84-02.04.2013.A429; AZ81-0204.2018.A369).

### Generation of Prm1-deficient mice

Single guide RNAs (sg1_ts: 5’-CACCGCGAAGATGTCGCAGACGG; sg1_bs: 5’- AAACCCGTCTGCGACATCTTCGC; sg2_ts: 5’-CACCGTGTATGAGCGGCGGCGA, sg2_bs: 5’-AAACTCGCCGCCGCTCATACAC) were tested in ES cells as described before [32]. Guides targeted exon 1 and exon 2 of *Prm1*.

CRISPR-Cas9-mediated gene editing of zygotes was performed as described before [32]. In brief, 6-8 weeks old B6D2F1 females were superovulated by intraperitoneal injections of 5 i.u. pregnant mare’s serum (PMS) and 5 i.u. human chorionic gonadotropin (hCG). Females were mated with B6D2F1 males and zygotes were isolated 0.5 dpc. Single guide RNAs (50 ng/µl each) were microinjected together with Cas9 mRNA (100 ng/µl). After culturing in KSOM medium for three days, developing blastocysts were transferred into the uteri of pseudo-pregnant CB6F1 foster mice. Offspring was genotyped by PCR and sequenced to identify founder animals. After first backcrossing to C57BL/6J mice, the F1 generation was sequenced. The allele (NM_013637.5:c.51_125del) was further back-crossed to C57BL/6J mice. Starting from the N3 generation analyses were performed, using male mice aged between 8-13 weeks.

### Prm2-deficient mice

*Prm2*-deficient mice (MGI: 5760133; 5770554) generated and analyzed by Schneider *et al*. [32, 33] were used for comparison.

### Genotyping and sequencing of mice

Primers flanking the gene edited region (Prm1_fwd: 5’-CCACAGCCCACAAAATTCCAC, Prm1_rev: 5’-TCGGACGGTGGCATTTTTCA) were used to amplify both the WT and edited allele (Cycling conditions: 2 min 95°C; 30x (30 sec 95°C; 30 sec 64°C; 35 sec 72°C); 5 min 72°C). PCR products (WT allele: 437 bp, *Prm1Δ*167: 270 bp) were separated on agarose gels.

PCR products were cloned using the TOPO™ TA Cloning™ Kit with pCR™2.1-TOPO™ (Thermo Fisher) according to the manufacturer’s instructions. Plasmids were transformed into E.cloni ® 10G Chemically Competent Cells (Lucigen, Middleton, WI, USA) according to the manufacturer’s instructions, isolated by alkaline lysis and sequenced by GATC/Eurofins (Cologne, Germany).

### Fertility assessment

Fertility was tested by mating male mice 1:1/1:2 to C57BL/6J females. Females were examined for presence of a vaginal plug daily. Plug positive females were separated and monitored. Pregnancies and litter sizes were recorded. A minimum of five plugs per male were evaluated.

### Immunohistochemistry (IHC)/ Immunofluorescence (IF)

Tissues were fixed in Bouin’s solution or paraformaldehyde (PFA) (4°C, overnight) processed in paraffin and 3 µm sections were generated. After deparaffinization, slides were treated with decondensation buffer, as described [33]. Heat mediated antigen retrieval was performed (citrate buffer pH 6.0) for 20 min, followed by blocking in Tris-HCl buffer (pH 7.4, 5% Bovine serum albumin, 0.5% Triton X-100) and primary antibody treatment overnight at 4°C. For IHC staining against protamines (anti-PRM1 (Hup1N) and anti-PRM2 (Hup2B) Briar Patch Biosciences, Livermore, CA, USA; 1:200), slides were treated with 3 % H_2_O_2_ for 30 min after decondensation. Biotinylated goat-anti-mouse (Dako, Glostrup, Denmark; E0433; 1:200) was used as secondary antibody (1 h, RT), processed using Vectastain Elite ABC-HRP Kit (Vector Laboratories, Burlingame, CA, USA; PK-6100) and stained with AEC-solution (Dako, AEC+ Substrate, K3469). Counterstain was performed using hematoxylin. For IF against 8-OHdG (Santa Cruz Biotechnology, Dallas, TX, USA; sc-66036; 1:200), goat-anti-mouse Alexa Fluor 488 (Thermo Fisher; A-11001; 1:500) was used as secondary antibody for 2 h at room temperature. Nuclei were stained using 1 µg/ml Hoechst (Thermo Fisher; 33342). 8-OHdG positive sperm were quantified using the Photoshop® counting tool. Two tubuli cross-sections per organ per mouse for three animals per genotype were analyzed.

### Macroscopic analysis of testis

Sections of Bouin-fixed testis were deparaffinized, hydrated, stained with Hemalum solution acid (Mayer) and Eosin Y solution (Carl Roth, Karlsruhe, Germany), dehydrated and mounted with Entellan® (Sigma-Aldrich/Merck, Darmstadt, Germany). Tubule diameters were determined measuring the horizontal and vertical diameters of at least 25 tubuli per testis cross-section. The number of elongated spermatids per tubules for a minimum of 5 tubules per mouse was counted with the ImageJ cell counter.

### Periodic Acid Schiff (PAS) Staining

PAS staining was performed as described [33]. After deparaffinization and re-hydration slides were incubated for 10 min in periodic acid (0.5%), rinsed in H_2_O, incubated 20 min with Schiff reagent, counterstained and mounted.

### Isolation of epididymal sperm

Sperm were isolated from the cauda epididymis by swim-out as described [32]. The epididymal tissue was incised multiple times and incubated in M2 medium (Sigma) or PBS at 37°C for 15-30 min.

### Transmission electron microscopy

Isolated sperm were pelleted (10,000 g, 2 min), fixed in 3% glutaraldehyde at 4°C overnight, washed with 0.1 M cacodylate buffer (2x 15 min), post-fixed with 2 % osmium tetroxide at 4°C for 2 h and again washed. After dehydration in an ascending ethanol series and contrasting in 70% (v/v) ethanol 0.5% (m/v) uranyl acetate (1 – 1.5 h, 4°C), samples were washed with propylenoxide (3x 10 min, RT) and stored in propylenoxide:Epon C (1:1, (v/v)) at 4°C overnight. Next, the pellets were embedded in Epon C (70 °C, 48 h). Ultra-thin sections were examined with transmission electron microscope CM10 equipped with analySiS imaging software. Using ImageJ, 100 sperm per sample were analyzed to determine the difference between the minimum and maximum grey value. Chromatin condensation status was categorized according to high (<150), intermediate (150-180) and low (>180) difference in grey scale.

### Assessment of sperm DNA integrity

Sperm genomic DNA was isolated as described [34] with minor adjustments. Briefly, sperm were incubated in 500 µl lysis buffer (1 M Tris-HCl pH 8.0, 3 M NaCl, 0.5 M EDTA, 20% (m/v) SDS) supplemented with 21 µl 1 M DTT, 2.5 µl 0.5% Triton-X100 and 40 µl 10 mg/ml proteinase K at 50°C overnight. After centrifugation (15,500 x g, 10 min), 1 µl 20 mg/ml glycogen and 1/10 vol 3 M NaAc were added to the supernatant. Precipitation was performed using absolute ethanol for 2 h at -80°C followed by 45 min at -20°C. The pellet was washed with 75% EtOH and dried in a Speed Vac DNA110 (Savant, Farmingdale, USA). DNA was dissolved in 30 µl TE buffer.

### Chromomycin A3 (CMA3) staining

Epididymal sperm were fixed in Carnoys solution (3:1 methanol:acetic acid, (v/v)), spread on microscopic slides and covered with 100 µl CMA3 solution (0.25 mg/ml CMA3 in Mcllvaine buffer (pH 7.0, containing 10 mM MgCl_2_)). After incubation for 20 min in the dark, slides were rinsed with Mcllvaine buffer and mounted with ROTI® Mount FluorCare DAPI (Carl Roth, Germany). 400 sperm per mouse were analyzed.

### Analysis of sperm membrane integrity

#### Eosin-Nigrosin staining

50 μl of sperm swim-out and 50 μl Eosin-Nigrosin stain (0.67 g eosin Y (color index 45380), 0.9 g sodium chloride, 10 g nigrosin (color index 50420), 100 ml ddH_2_O) were mixed and incubated for 30 sec. 30 μl of the mix was pipetted onto microscope slides, smeared and mounted with Entellan® (Merck, Darmstadt, Germany). 200 sperm per animal were analyzed.

#### Hypoosmotic swelling test

100 μl of sperm swim-out was mixed with 1 ml pre-warmed HOS solution (1.375 g D-fructose, 0.75 g sodium citrate dihydrate, 100 ml ddH_2_O) and incubated for 30 min at 37°C. The solution was dropped onto a microscopic slide, covered with a cover slip and analyzed within 1 h. 200 sperm per animal were evaluated.

#### RNA sequencing (RNAseq) and differential expression analysis

RNA was extracted from whole testis of three individuals per genotype. After removal of the tunica albuginea, testes were homogenized in TRIzol™ and processed according to the manufacturers protocol (Thermo Fisher). RNA integrity (RIN) was determined using the RNA Nano 6000 Assay Kit with the Agilent Bioanalyzer 2100 system (Agilent Technologies, Santa Clara, CA, USA). RIN values were > 7 for all samples. RNA sample quality control, library preparation (QuantSeq 3’-mRNA Library Prep (Lexogen, Greenland, NH, USA)) and RNAseq were performed by the University of Bonn Core facility for Next Generation Sequencing (NGS). Sequencing was performed on the Illumina HiSeq 2500 V4 platform, producing >10 million, 50bp 3’-end reads per sample.

Samples were mapped to the mouse genome (GRCm38.89) using HISAT2 2.1 [35] and transcripts were quantified and annotated using StringTie 1.3.3 [36]. Gene annotation was retrieved from the Ensembl FTP server (ftp://ftp.ensembl.org)(GRCm38.89). The python script (preDE.py) included in the StringTie package was used to prepare DEseq2-compatible gene-level count matrices for analysis of differential gene expression. Mapping to the *Prm1* genomic location was visualized using the Integrative Genomics Viewer (IGV) [37].

Differential expression was analyzed using DESeq2 1.16.1 [38]. The adjusted p-value (Benjamini-Hochberg method) cutoff for DE was set at < 0.05, log2 fold change of expression (LFC) cutoff was set at > 1. We performed GO term and pathway overrepresentation analyses on relevant lists of genes using the PANTHER gene list analysis tool with Fisher’s exact test and FDR correction [39].

#### Mass spectrometry and differential protein abundance analysis

Sperm basic nuclear proteins from three WT, *Prm1-/-* and *Prm2-/-* mice were isolated as described below and used for mass spectrometric analysis. Peptide preparation, LC-MS and differential abundance (DA) analysis were performed at the University of Bonn Core facility Mass Spectrometry.

Peptide preparation: Protein solutions (5.5 M urea, 20% 2-mercaptoethanol, 5% acetic acid) were dried in a vacuum concentrator and subjected to in solution preparation of peptides as described previously [40]. Briefly, cysteines were alkylated with acrylamide and digested with trypsin, followed by desalting.

LC-MS measurements were performed according to Arévalo *et al*. [40]. Briefly, peptides were separated on a self-packed reversed-phase column within a 90 min gradient. Peptide ions were analyzed with an Orbitrap Lumos mass spectrometer in data-dependent mode with a top-speed method. Precursors and fragment ions were recorded with the Orbitrap detector. Raw data processing and was performed with Proteome Discoverer software in combination with Mascot server version 2.6.1 using *Mus musculus* sequences from SwissProt (2021/03, including isoforms), and contaminants (cRAP, [41]). Mascot results were filtered for 1% FDR on the basis of q-values from the percolator algorithm [42]. Spectra with identifications below 1% q-value were sent to a second round of database search with semi-tryptic enzyme specificity Summed abundances were used for relative quantification.

Differential abundance (DA) analysis: DA analysis was performeds using the Bioconductor package proDA [43] using peptide spectrum matches (PSM) level data extracted from Protein Discoverer. Only proteins detected in all genotypes and all replicates with more than two peptides were included in the analysis. The data were log2 transformed and median normalized prior to DA analysis to ensure comparability. The proDA package is based on linear models and utilized Bayesian priors to increase power for differential abundance detection [43]. Proteins with a log2 fold change (LFC) of >1 and false discovery rate adjusted p-value (FDR) <0.05 were considered differentially abundant compared to the WT. Plots were generated using the R-package ggplot2 [44].

#### Sperm nuclear morphology analysis

Epididymal sperm were analyzed using the ImageJ plugin “Nuclear_Morphology_Analysis_1.18.1_standalone” [45] as described previously [33]. In brief, sperm were fixed in Carnoys solution (3:1 methanol: acetic acid (v/v)), spread on slides, mounted with ROTI® Mount FluorCare DAPI (Carl Roth, Karlsruhe, Germany) and imaged at 100-fold magnification. A minimum of 100 sperm heads per sample from four biological replicates were analyzed.

#### Sperm motility analysis

Epididymal sperm swim out was performed in 1 ml sterile filtered THY medium (138 mM NaCl, 4.8 mM KCl, 2 mM CaCl_2_, 1.2 mM KH_2_PO_4_, 1 mM MgSO_4_, 5.6 mM glucose, 10 mM HEPES,

0.5 mM sodium pyruvate, 10 mM L-lactate, pH 7.4, 310-320 mOsm) for 15 min at 37°C. Next, sperm were diluted 1:20 – 1:50 in dilution medium (3 mg/ml BSA in THY medium). 30 µl of dilution were pipetted onto a glass slide equipped with a spacer and cover slip, placed on a heated slide holder (37°C) and analyzed under an inverted microscope (Leica, Wetzlar, Germany) equipped with a camera (acA1920-155ucMED; Basler AG, Ahrensburg, Germany). The movement of sperm was recorded at 100 frames/sec for 3 sec and analyzed in ImageJ. The produced “z project” was used to distinguish and count moving and non-moving sperm (n = 100 sperm/ mouse).

#### Analysis of sperm basic nuclear proteins

Isolation of sperm nuclear proteins was performed according to Soler-Ventura *et al*. [46]. Briefly, sperm were counted, washed in PBS, pelleted and resuspended in 200 µl buffer containing 4 µl 1 M Tris pH8, 0.8 µl 0.5 M MgCl_2_ and 5 µl Triton X-100. After centrifugation the pellet was mixed with 1 mM PMSF. Lysed cells were mixed with solutions containing PMSF, EDTA, DTT, GuHCl and vinylpyridine and incubated for 30 min at 37°C. Addition of EtOH precipitates DNA. Proteins are dissolved in 0.5 M HCl and precipitated with TCA. After acetone washes the proteins are lyophilized and resuspended in sample buffer (5.5 M Urea, 20% 2-mercaptoethanol, 5% acetic acid).

Next, the nuclear proteins were separated on a pre-electrophorized 15% acid-urea polyacrylamide gel (2.5 M urea, 0.9 M acetic acid, and 15% acrylamide/ 0.1% N,N′-Methylene bis-acrylamide, TEMED and APS) and visualized with Coomassie Brilliant Blue. Quantification was performed utilizing ImageJ as described previously [40].

#### Statistics

Values are, if not indicated otherwise, presented as mean values with standard deviation. Statistical significance was calculated by two-tailed, unpaired Student’s t-test and a value of p < 0.05 was considered significant (p < 0.05= *; p < 0.005= **; p < 0.001= ***).

## Results

### CRISPR-Cas9-mediated gene editing produces Prm1-deficient mice

*Prm1*-deficient mice were generated using CRISPR-Cas9-mediated gene editing. Guide RNAs targeting exon 1 and exon 2 of *Prm1* and Cas9 mRNA were injected into zygotes. From the 13 pups obtained four contained a deletion in the *Prm1* coding region. Those animals were mated to C57BL/6J mice and from the offspring the *Prm1* locus was sequenced. We selected a mouse carrying a 167 bp in frame deletion in the *Prm1* coding region (**Fig. 1a**) and established a PCR-based genotyping (**Fig. 1b**).

**Fig. 1.**
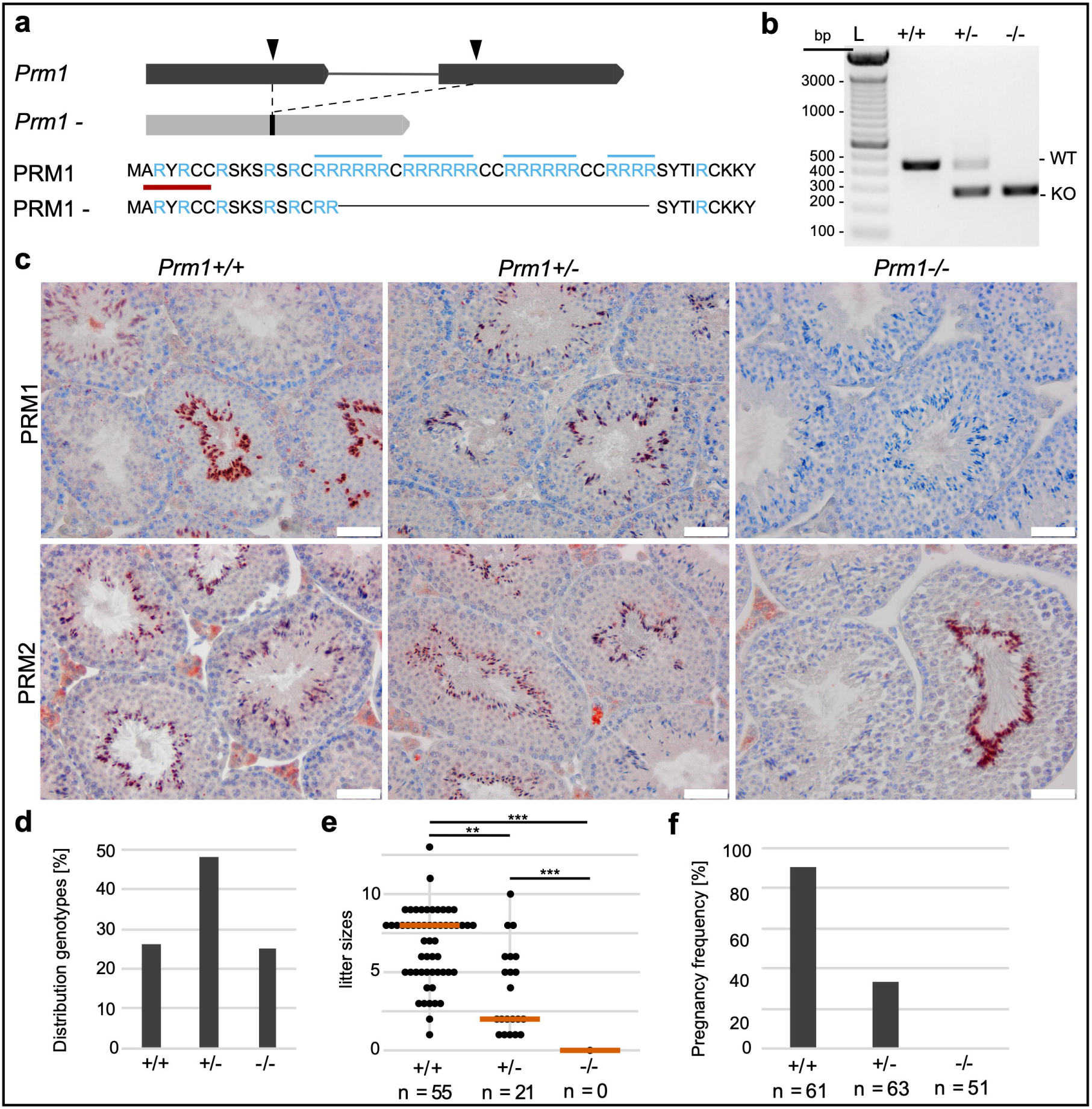
Establishment of *Prm1*-deficient mice and fertility analysis. **(a)** Graphical representation of CRISPR-Cas9-mediated gene editing of the *Prm1* locus. Two guide RNAs were used (indicated by black arrow heads); targeting the *Prm1* coding sequence in exon 1 and exon 2, respectively. A 167 bp in-frame deletion was generated, leading to loss of crucial arginine-rich DNA binding sites (marked in blue). The epitope of the anti-PRM1 antibody used in Fig.1c is marked in red. **(b)** Agarose gel of genotyping polymerase chain reaction of *Prm1+/+, Prm1+/-* and *Prm1-/-* mice. Amplification of the wild type *Prm1* or the *Prm1-*allele generates products of 437 bp or 270 bp, respectively. L = ladder **(c)** Immunohistochemical staining against PRM1 and PRM2 on Bouin-fixed, paraffin-embedded testis sections of *Prm1+/+, Prm1+/-* and *Prm1-/-* mice counterstained with hematoxylin. Scale: 50 µm **(d)** Mendelian distribution of genotypes (n = 10 litters) from crossings of *Prm1+/-* males and females. **(e)** Scatter plot of litter sizes monitored after mating with female WT C57BL/6J mice. n = number of pregnancies produced by 12 *Prm1+/+*, 9 *Prm1+/-* and 9 *Prm1-/-* males, respectively. The mean litter size is indicated by vermillion lines. **(f)** Pregnancy frequency (%) after mating with female WT C57BL/6J mice. n = number of plugs produced by 12 *Prm1+/+*, 9 *Prm1+/-* and 9 *Prm1-/-* males, respectively.

In order to validate the deletion, 3’-mRNA sequencing of whole testis of WT (*Prm1+/+*) and *Prm1-/-* males was performed. In *Prm1-/-* males the transcripts mapped to the 5’ and the 3’and ends of the *Prm1* locus while we could not detect transcripts from the central, deleted area, which encodes for crucial arginine sites required for DNA binding (**Fig. 1a, Supplementary Fig. 1**).

Next, we used immunohistochemical (IHC) staining with an anti-PRM1 antibody, targeting an epitope at the N-terminus of PRM1 (marked in red (**Fig. 1a**)) in order to determine, whether the potential transcripts of the gene-edited allele result in the production of a truncated PRM1 protein. However, we could not detect a signal in testis sections of *Prm1-/-* males (**Fig. 1c**). This strongly suggests nonsense mediated RNA decay of the potential transcript and demonstrates, that the deletion introduced by CRISPR-Cas9 results in a functional *Prm1* null allele. PRM1 was detected in elongating spermatids and spermatozoa in wildtype (*Prm1+/+*) and *Prm1+/-* testis sections. PRM2 was present in all genotypes.

Mating of *Prm1+/-* males with *Prm1+/-* females produced approximately 50% *Prm1+/-* and 25% *Prm1+/+* or *Prm1-/-* animals respectively (**Fig. 1d**), suggesting that the deletion did not interfere with embryonic development.

### Prm1-/- male mice are infertile, while Prm1+/- are subfertile

After establishing and validating the *Prm1*-deficient line, we performed fertility tests with *Prm1+/-* and *Prm1-/-* males. *Prm1+/-* males are subfertile, while *Prm1-/-* males are sterile (**Fig. 1e**). None of the nine *Prm1-/-* males tested was able to generate offspring. *Prm1+/-* males generate smaller average litter sizes (mean: 3.81 ± 2.75) compared to WT males (mean: 6.75 ± 2.39) (**Fig. 1e**). Additionally, the pregnancy frequency of *Prm1+/-* males is significantly reduced (**Fig. 1f**). Only about 33% of the monitored copulations with *Prm1+/-* males resulted in pregnancies. These results indicate that loss of one allele of *Prm1* already reduces male mice fecundity.

### Spermatogenesis unaffected in Prm1-deficient mice

In order to test, whether the deletion of *Prm1* affects spermatogenesis, we analyzed standard male fertility parameters. The relative testis mass (**Fig. 2a**), the average seminiferous tubuli diameter (**Fig. 2b**) and the number of elongating spermatids per seminiferous tubule cross section (**Fig. 2c**) are not reduced in *Prm1+/-* or *Prm1-/-* animals when compared with *Prm1+/+* animals. Spermatozoa lining up at the lumen of stage VII-VIII seminiferous tubules can be detected in *Prm1*-deficient mice (**Fig. 2d**). Spermatids undergo differentiation, elongate and acrosomal structures and flagellae are formed in *Prm1-*deficient mice (**Fig. 2e**). These results suggest that spermatogenesis is unaffected in *Prm1+/-* and *Prm1-/-* mice.

**Fig. 2.**
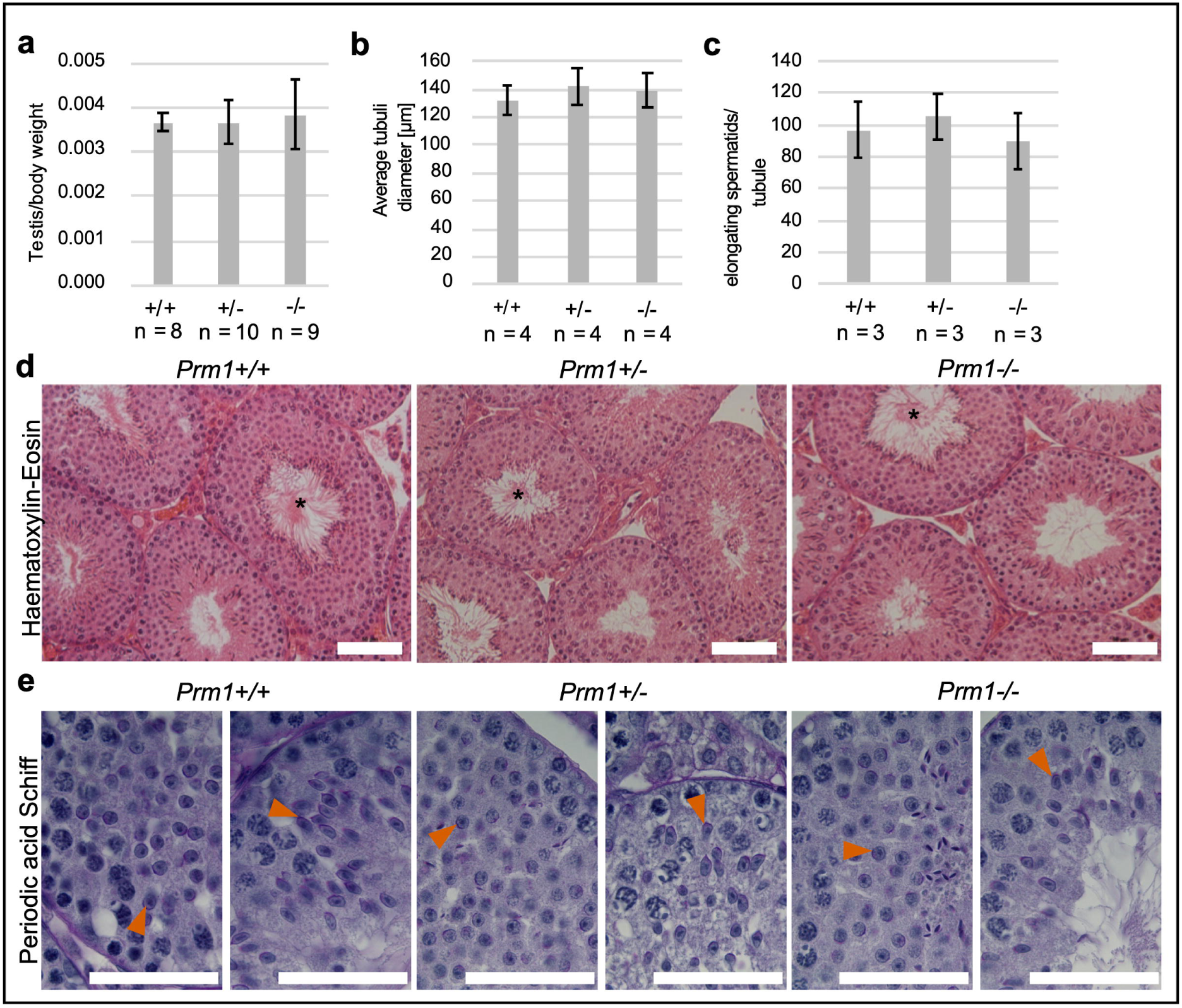
Spermatogenesis of *Prm1*-deficient mice. **(a)** Testis to body weight ratio of *Prm1+/+, Prm1+/-* and *Prm1-/-* males (n = 8-10). **(b)** Average diameter of seminiferous tubules of *Prm1+/+, Prm1+/-* and *Prm1-/-* mice (n = 4). 25 tubules per mouse were evaluated. **(c)** Quantification of elongating spermatids per seminiferous tubule cross-section in *Prm1+/+, Prm1+/-* and *Prm1-/-* males (n = 3). 5 tubules per mouse were evaluated. **(d)** Hematoxylin-Eosin staining of testis of *Prm1+/+, Prm1+/-* and *Prm1-/-* males. Tubules at stage VII-VIII of the epithelial cycle with spermatozoa lining up at the edge of tubule lumen are marked with asterisks. Scale: 50 µm **(e)** Periodic acid Schiff staining of testis of *Prm1+/+, Prm1+/-* and *Prm1-/-* males. Acrosomal structures are indicated by vermillion arrow heads. Scale: 50 µm

### Epididymal Prm1-deficient sperm display ROS-mediated DNA damage

Since PRM1 is necessary for DNA hypercondensation, we evaluated chromatin compaction of epididymal sperm. Transmission electron micrographs of epididymal sperm revealed defects in chromatin hypercondensation in *Prm1-/-* sperm compared to *Prm1+/-* and *Prm1+/+* sperm (**Fig. 3a**). While approximately 80 - 85% of *Prm1+/-* and *Prm1+/+* epididymal sperm nuclei appear electron dense indicative for condensed chromatin, only around 29% of *Prm1- /-* sperm nuclei seem fully condensed (**Fig. 3b**). Additionally, epididymal sperm from *Prm1-/- mice* present with membrane damage and disrupted acrosomes in transmission electron micrographs (**Fig. 3c**).

**Fig. 3.**
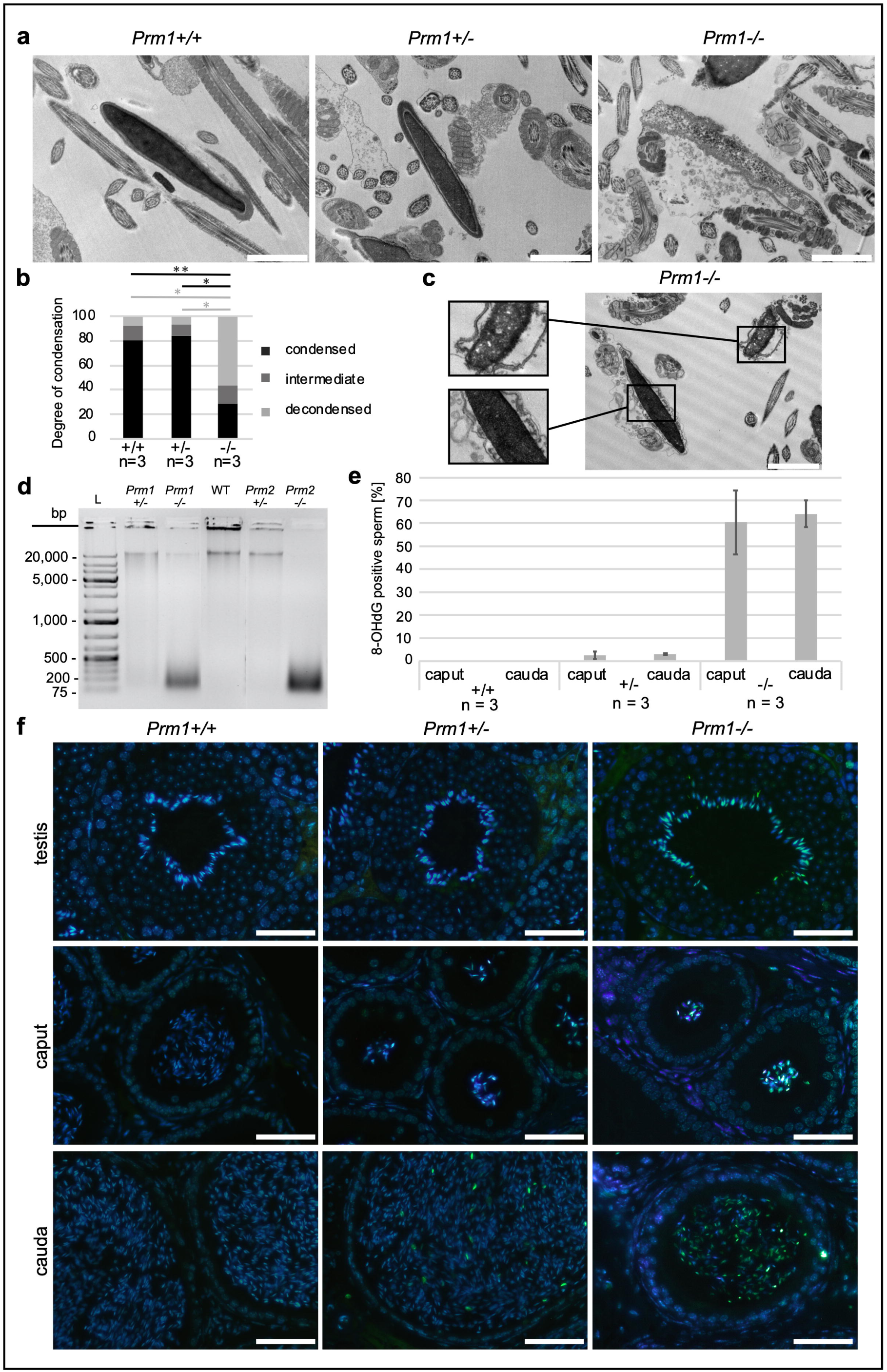
Analysis of chromatin condensation and ROS-induced DNA damage in epididymal *Prm1*-deficient sperm. **(a)** Representative transmission electron micrographs of *Prm1+/+, Prm1+/-* and *Prm1-/-* epididymal sperm. Scale: 2 µm **(b)** Quantification of DNA condensation of epididymal sperm from *Prm1+/+, Prm1+/-* and *Prm1-/-* males (n = 3). 100 sperm per male were analyzed. **(c)** Transmission electron micrograph of *Prm1-/-* epididymal sperm. Scale: 2 µm **(d)** Agarose gel loaded with genomic DNA isolated from epididymal sperm of *Prm1+/-, Prm1-/-, Prm2+/-, Prm2-/-* and WT males separated by electrophoresis. L = ladder **(e)** Percentage of 8-OHdG positive sperm on tissue sections of caput and cauda epididymis of *Prm1+/+, Prm1+/-* and *Prm1-/-* mice (n =3). **(f)** Representative immunofluorescent staining against 8-OHdG in testis, caput epididymis and cauda epididymis tissue sections from *Prm1+/+, Prm1+/-* and *Prm1-/-* males. Scale: 50 µm

To assess DNA damage, genomic DNA isolated from epididymal sperm was separated by agarose gel electrophoresis. DNA from WT sperm presents as a single band of high molecular weight indicative for intact DNA. Contrary, the majority of DNA isolated from *Prm1-/-* epididymal sperm is detected as fragments of approximately 100-500 bp indicative of strong DNA degradation. While DNA of sperm from *Prm2-/-* male mice is completely fragmented, a small proportion of DNA in *Prm1-/-* sperm is presented as a high molecular weight band indicating that a small portion of DNA from *Prm1*-/- sperm remains intact. DNA from *Prm1+/-* sperm displays a weak smear indicative for low, but detectable level of DNA-degradation (**Fig. 3d**). This suggests that loss of one *Prm1* allele leads to low levels of DNA damage. This is in contrast to *Prm2*, where loss of one allele was tolerated and DNA did not show any sign of degradation.

Since similar DNA damage have been described for *Prm2*-/- sperm and have been correlated to increased reactive oxygen species (ROS) levels during epididymal transit [32, 33], we stained testicular and epididymal tissue sections for 8-OHdG (8-hydroxydeoxyguanosine), a marker for oxidative stress induced DNA lesions. In tissue sections from epididymides of *Prm1-/-* mice, 60% of caput sperm and 64% of cauda sperm stained 8-OHdG-positive (**Fig. 3e-f**). In epididymides of *Prm1+/-* mice a small number of sperm stained positive for 8-OHdG (mean: 2.6% in caput and 3.0% in cauda epididymis). In contrast, on sections of *Prm1+/+* mice, no staining was detected. This shows, that the low level of DNA damage detected in *Prm1+/-* males is most likely restricted to the few 8-OHdG positive sperm and not due to low level of DNA damage in all sperm. Of note, the majority of the sperm from *Prm1-/-* mice stain 8-OHdG-positive in the testis (**Fig. 3f**).

### Epididymal Prm1-deficient sperm display impaired membrane integrity, nuclear head morphology changes and sperm motility defects

To characterize possible secondary effects of ROS, we next used Eosin-Nigrosin staining and a hypoosmotic swelling test to test for sperm membrane integrity. *Prm1-/-* epididymal sperm display severe membrane damage indicative of inviable sperm, while no significant difference between *Prm1+/-* and *Prm1+/+* sperm was detected (**Fig. 4a, b, Supplementary Fig. 2a, b**).

**Fig. 4.**
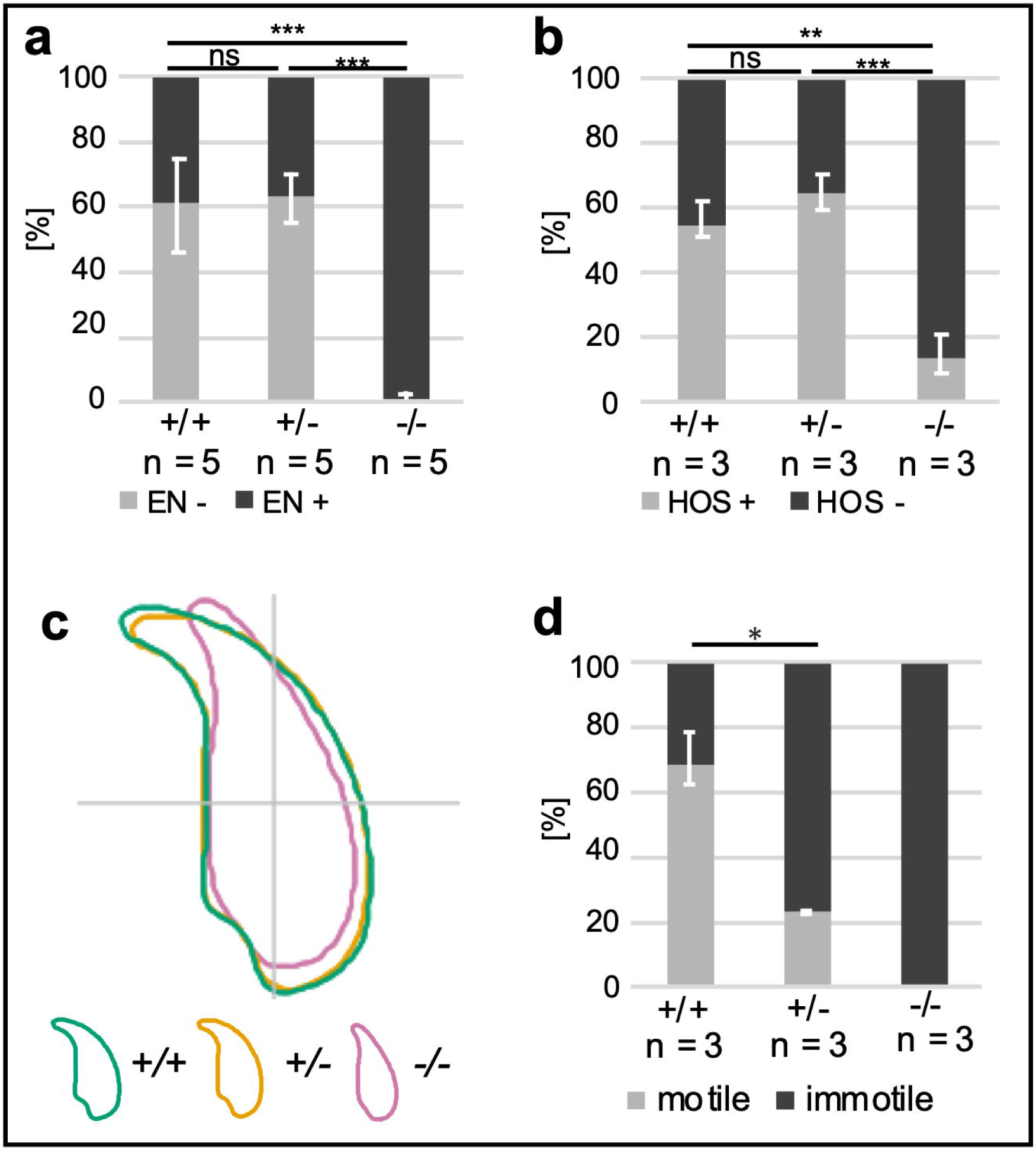
Secondary effects on *Prm1*-deficient epididymal sperm. **(a)** Eosin-Nigrosin (EN) staining: Quantification of EN positive and EN negative sperm (%) from *Prm1+/+, Prm1+/-* and *Prm1-/-* males. (n = 5) **(b)** Hyperosmotic swelling test: Quantification of HOS positive and HOS negative sperm (%) from *Prm1+/+, Prm1+/-* and *Prm1-/-* males (n =3). **(c)** Nuclear head morphology analysis for *Prm1+/+, Prm1+/-* and *Prm1-/-* sperm. Consensus shapes of sperm heads are depicted. 4 males per genotype and a minimum of 100 sperm per animal were analyzed. **(d)** Quantification of motile and immotile sperm (%) from *Prm1+/+, Prm1+/-* and *Prm1-/-* males (n = 3).

For analysis of epididymal sperm head morphology we used a high-throughput ImageJ plugin [45] and generated a consensus shape visualizing the overall head shape of the population analyzed. *Prm1-/-* sperm lose the typical hooked sperm head shape (**Fig. 4c, Supplementary Fig. 3a**). While the head shape of *Prm1-/-* sperm displays higher variability (**Supplementary Fig. 3b**), they appear smaller with a mean area of 14.92 µm^2^ (95% CI 14.92 ± 0.26) compared to 19.82 µm^2^ (95% CI 19.82 ± 0.10) and 19.47 µm^2^ (95% CI 19.47 ± 0.13) for *Prm1+/+* and *Prm1+/-* sperm heads, respectively (**Supplementary Fig. 3c**). Further, *Prm1-/-* sperm heads are more elliptic (**Supplementary Fig. 3d**) and thinner (**Supplementary Fig. 3e**). *Prm1+/-* sperm heads show a slightly stronger hook curvature resulting in a reduced maximum ferret of 8.07 µm (95%b CI 8.07 ± 0.04) compared to 8.38 µm (95% CI 8.38 ± 0.04) for *Prm1+/+* sperm (**Supplementary Fig. 3f**). The reduction in maximum ferret is significant, however, should be interpreted carefully, when considering the general variability but clear overlap in sperm head shapes depicted for *Prm1+/-* and *Prm1+/+* sperm populations (**Supplementary Fig. 3b**). These results suggest, that loss of one allele of *Prm1* does not affect sperm head shape dramatically.

Next, we analyzed the percentage of motile sperm isolated from the cauda epididymis (**Fig. 4d**). Strikingly, *Prm1+/-* sperm showed a marked reduction in sperm motility. Only around 23% of the *Prm1+/-* sperm were motile. In contrast, 77% of WT sperm were motile, while *Prm1*-deficient sperm are completely immotile. So, the reduction in motility contributes to the sub/infertility seen.

### Transcriptional and proteomic profiling reveals differences in Prm1 and Prm2 deficient males

To address the question, whether transcriptional silencing is affected upon loss of protamines, we performed transcriptomic and proteomic analyses. 3’-mRNA sequencing of the whole testis revealed that in *Prm1-/-* testis 99 genes are higher and 11 lower expressed, while in *Prm1+/-* testis 28 genes were higher and 39 were lower expressed, both compared to WT testis (**Fig. 5a, Supplementary Material 1**). In *Prm1-/-* testis pathway enrichment for immune related genes (*Il1b, Ccl5, Saa3, Atp6ap1, Rsad2, Cxcl10, Ifit1, Mmp13, Clec4e, Zghhc*) was identified. These transcripts were slightly higher abundant in *Prm1-/-* testis compared to WT testis, but showed low levels of expression (**Supplementary Material 1**). This might either indicate a reaction to ROS-mediated damage of the sperm in testis or an unspecific failure in transcriptional silencing.

**Fig. 5.**
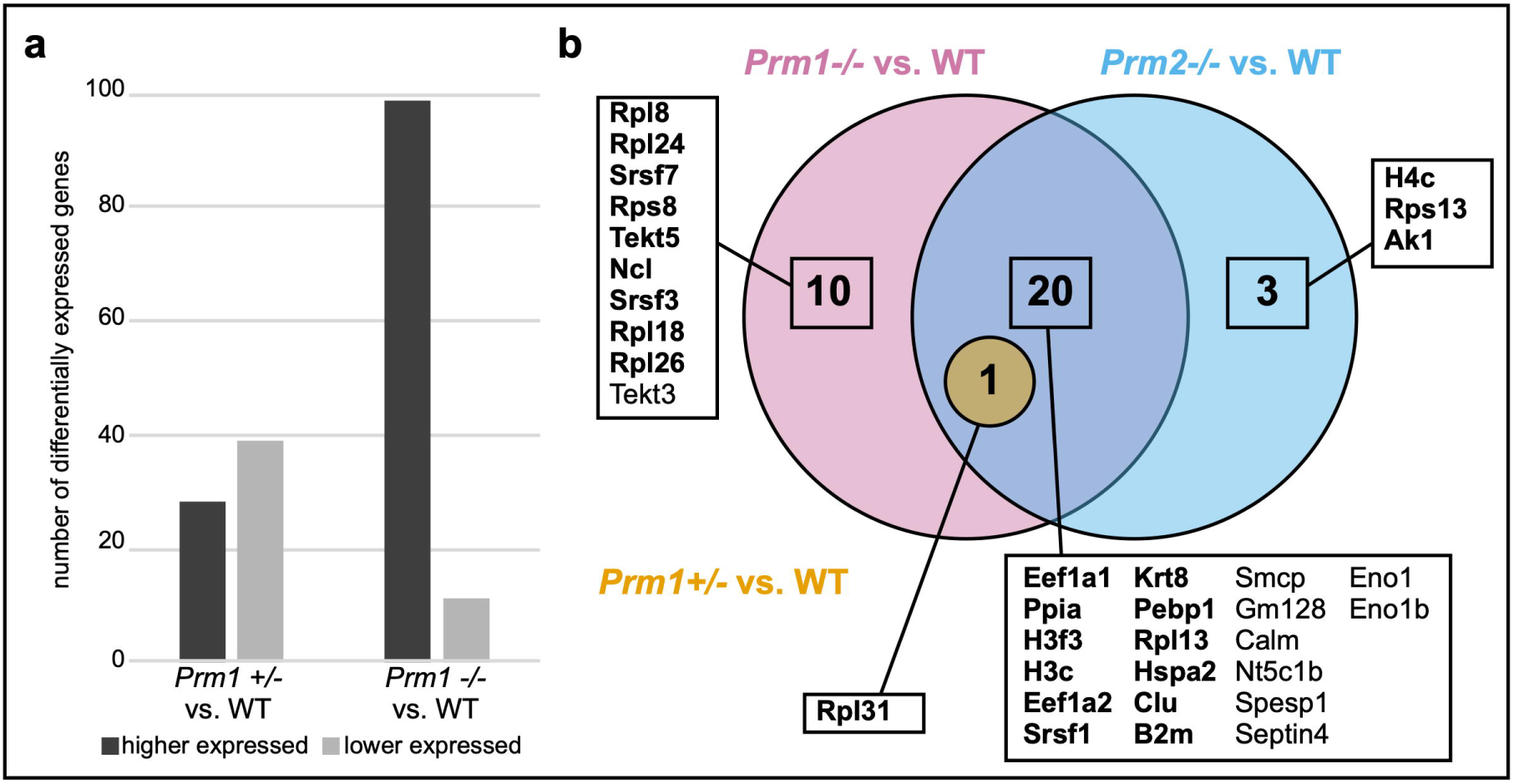
Differentially expressed genes in the testis and altered protein abundances in sperm in protamine-deficient males. **(a)** Number of differentially expressed genes subdivided into higher and lower expressed genes in testis of *Prm1+/-* and *Prm1-/-* males compared to WT males, respectively. **(b)** Venn diagram illustrating changes in abundances of proteins from sperm nuclear protein extractions of *Prm1-/-, Prm1+/-* and *Prm2-/-* males compared to WT, respectively. Proteins that were higher abundant are depicted in bold letters. Non-bold proteins were lower abundant compared to WT.

In order to determine whether proteins might be differentially abundant in mature sperm, we analyzed basic nuclear protein extracts of *Prm1+/-, Prm1-*/-, *Prm2-/-* and WT sperm with MassSpec. In *Prm1-/-* samples 31 proteins were differentially abundant compared to WT sperm (**Fig. 5b, Supplementary Material 2**). Of these, 21 were also differentially abundant in *Prm2-/-* samples. Proteins related to translation, mRNA splicing and protein folding (EEF1A1, EEF1A2, RPL13, RPL31, SRSF1 and PPIA) were detected to be higher abundant in *Prm1-/-* or *Prm2-*/- sperm compared to WT sperm. Additionally, histones (H3F3, H3C) were found to be higher abundant in *Prm1-/-* and *Prm2-/-* sperm, indicating increased H3 histone retention. In addition, in *Prm2-/-* males also histone H4C was higher abundant. In *Prm1-/-* samples, further proteins were detected to be higher abundant related to translation and mRNA splicing (RPL8, RPS8, RPL18, RPL24, RPL26, SRSF3, SRSF7). Proteins related to stress response and apoptosis (B2M, CLU, HSPA2) were also higher abundant in sperm lacking PRM1 or PRM2. This we expected to be a stress response due to the increased ROS-mediated sperm damage detected. SMCP and SPESP1, proteins important for sperm motility and sperm-egg fusion on the other hand are lower abundant in both *Prm1-/-* and *Prm2-/-* samples. Only one protein, the ribosomal protein RPL31, which was also identified in *Prm1-/-* and *Prm2-/-* samples, was higher abundant in *Prm1+/-* sperm nuclear extracts compared to WT sperm. The fact that there is only one non-protamine protein differentially abundant in *Prm1+/-* nuclear extracts, suggests that the *Prm1+/-* sperm protein profile is not causative of the subfertility observed.

### Protamine and nuclear protein content are altered in protamine deficient epididymal sperm

Next, we analyzed the level of protamination using Chromomycin A3 (CMA3), a dye competing with protamines to bind CG-rich regions to the minor groove of DNA [47]. While 98% of *Prm1+/-* sperm show CMA3 staining, only around 29% of *Prm2+/-* sperm showed a CMA3-signal (**Fig. 6a, b**). These data suggest that chromatin in *Prm1*+/- and *Prm2+/-* sperm is either not fully or not correctly protaminated, with the effects being more dramatic in *Prm1+/-* sperm. Of note, sperm from *Prm1-/-* and *Prm2-/-* mice could not be analyzed due to the fact that severe DNA fragmentation interfered with the staining procedure.

**Fig. 6.**
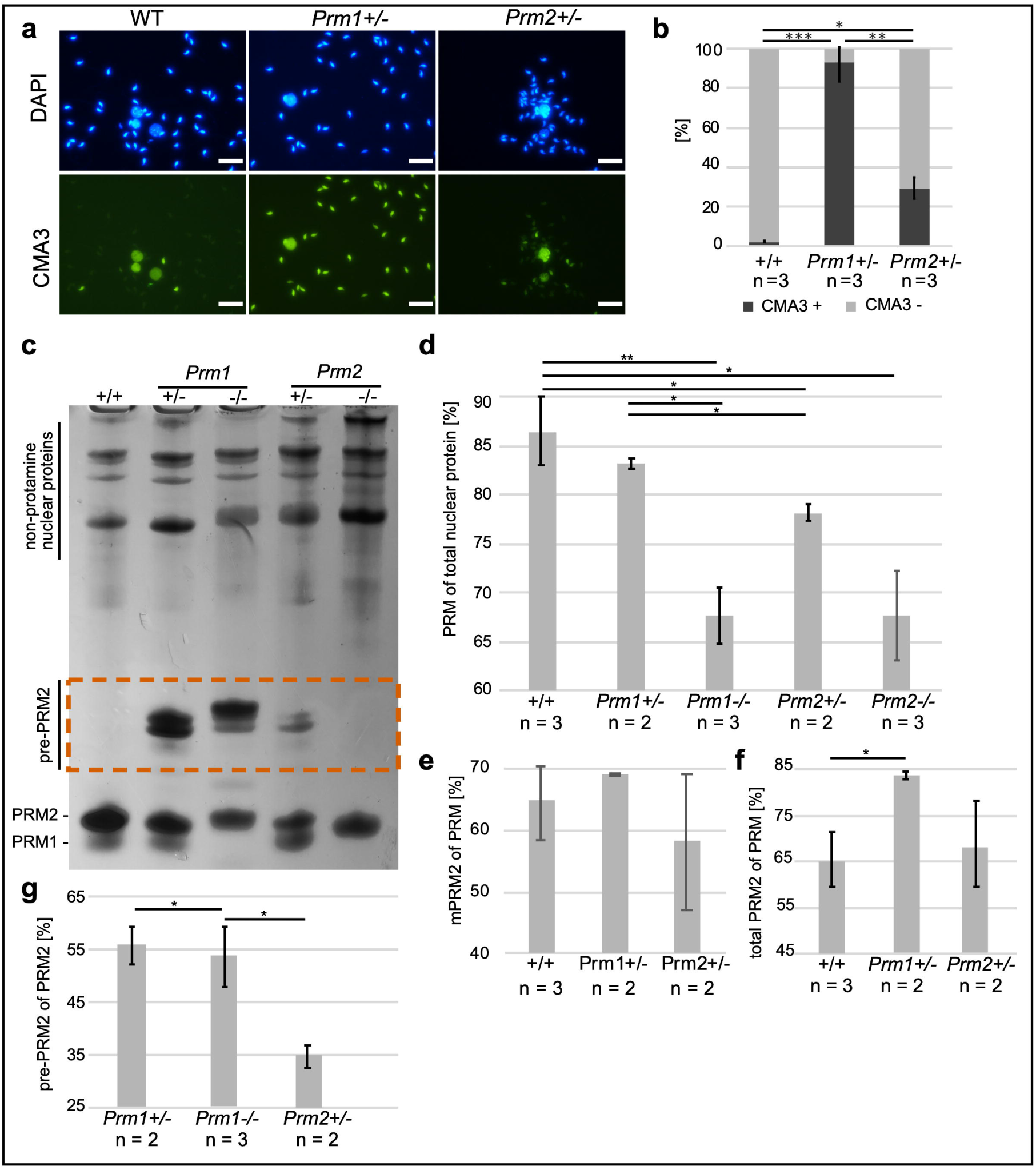
Sperm nuclear protein analysis in protamine-deficient sperm. **(a)** Representative pictures of CMA3 staining of *Prm1+/+, Prm1+/-* and *Prm2+/-* epididymal sperm heads taken at same exposure time. DAPI was used as counter stain. Scale: 20 µm. **(b)** Average percentage of CMA3 positive and negative sperm in *Prm1+/-, Prm2+/-* and WT males (n = 3). A minimum of 400 sperm per male were analyzed. (c) Representative acid-urea polyacrylamide gel (AU-PAGE) of nuclear protein extractions from WT, *Prm1+/-, Prm1-/-, Prm2+/-* and *Prm2-/-* epididymal sperm. Non-protamine nuclear proteins can be detected at the top of the AU-PAGE. PRM1 and PRM2 run at the bottom of the gel. PRM2 precursor forms (pre-PRM2) run higher than PRM (marked by vermillion box). **(d)** Percentage of PRM of total nuclear protein in nuclear protein extractions from WT, *Prm1+/-, Prm1-/-, Prm2+/-* and *Prm2-/-* epididymal sperm. **(e)** Percentage of mPRM2 of PRM in nuclear protein extractions from WT, *Prm1+/-* and *Prm2+/-* epididymal sperm. **(f)** Percentage of total PRM2 (including pre-PRM2) of PRM in nuclear protein extractions from WT, *Prm1+/-* and *Prm2+/-* epididymal sperm. **(g)** Percentage of pre-PRM2 of PRM2 in nuclear protein extractions from *Prm1+/-, Prm1-/-* and *Prm2+/-* epididymal sperm.

To further analyze the relative protamine content and protamination of epididymal sperm of *Prm1+/-, Prm1-/-, Prm2+/-*, Prm2-/- and WT mice in more detail, basic nuclear proteins were separated on acid-urea polyacrylamide gels (AU-PAGE). Most interestingly, in sperm from *Prm1+/-* and *Prm1-/-* mice PRM2 precursors (pre-PRM2) were detected suggesting disturbances in processing of PRM2 upon loss of PRM1 (**Fig. 6c**, vermillion box). Further, we quantified the relative amounts of nuclear proteins within individual samples (**Fig. 6d-g, Supplementary Fig. 3**).

In WT epididymal sperm protamines account for around 86% of the total nuclear proteins (**Fig. 6d**). Interestingly, while the difference in sperm protamine content in *Prm1+/-* is not significant (83%), the protamine content is significantly reduced in sperm from *Prm2+/-* (78%), *Prm1-/-* (67%) and *Prm2-/-* (67%) mice (**Fig. 6d**). These results might help explaining the increased histone retention in *Prm1-/-* and *Prm2-/-* sperm, as detected by MassSpec. While the relative amount of mPRM2 to total protamine is not significantly different in *Prm1+/-* and *Prm2+/-* sperm compared to WT (**Fig. 6e**), the total amount of PRM2 (mPRM2 + pre-PRM2) is significantly higher in *Prm1+/-* sperm (83%) only (**Fig. 6f**). Taking these data in account, in *Prm1+/-* sperm the PRM1:PRM2 ratio is shifted to approximately 1:5 while the species-specific protamine ratio of 1:2 is maintained in *Prm2+/-* sperm which is comparable to WT [14]. Consequently, the pre-PRM2 content of total PRM2 is significantly larger in *Prm1+/-* and *Prm1-/-* sperm compared to *Prm2+/-* sperm (**Fig. 6g**).

## Discussion

In this study, mice deficient for *Prm1* were generated using CRISPR-Cas9-mediated gene editing. *Prm1*-/- male mice are infertile, while loss of one allele of *Prm1* results in subfertility. *Prm1-/-* sperm show severe DNA fragmentation, high levels of 8-OHdG, destructed membranes and complete immotility. *Prm1+/-* sperm show moderate ROS-induced DNA damage, reduced sperm motility and a shifted PRM1:PRM2 ratio. *Prm1-/-* and *Prm1+/-* sperm contain high levels of incompletely processed PRM2 suggesting that PRM1 is necessary for correct PRM2 processing.

Protamine deficient mouse models have been described and associated with male factor infertility in previous studies [28-31]. Contrary to previous studies, we show that *Prm1*+/- males are able to produce offspring by natural breeding. *Prm1*-deficient chimeras, that have been generated by classical gene-targeting techniques, were reported to be sterile [28], excluding mouse line establishment and detailed studies on *Prm1*-deficiency. Takeda *et al*. were, however, able to generate viable offspring from *Prm1*+/- males by *in vitro* fertilization (IVF) of zona-free oocytes [30]. Further, Mashiko *et al*. reported that CRISPR-Cas9-mediated *Prm1+/-* mice are infertile, however detailed fertility statistics and phenotypical analysis of *Prm1*-deficient mice were not performed [31]. Since the *Prm1+/-* males produced by us are subfertile, we were able to generate and analyze *Prm1-/-* mice. Takeda *et al*. used a different mouse strain (C57BL/6J x DBA, backcrossed to CD1) and ES-targeting technology, which might explain the differences in *Prm1+/-* fertility. While Mashiko *et al*. used both the identical strain (C57BL/6J x DBA, backcrossed to C57BL/6J) and technology, they might not have performed a sufficiently exhaustive fertility analysis in order to detect subfertility.

Spermatogenesis seems unaffected in *Prm1*-/- (and *Prm1+/-*) mice compared to WT mice. Similar results were described for *Prm2*-/- mice, where spermatogenesis appears normal [32], epididymal sperm however show severe damage. In *Prm2-/-* mice it has been reported that an oxidative stress-mediated destruction cascade is initiated during epididymal sperm maturation [32, 33]. While it is well known that low levels of ROS are required for proper sperm function, high levels cause sperm pathologies [48]. Accumulation of ROS and loss of the antioxidant capacity of *Prm2*-/- sperm caused severe DNA fragmentation, sperm immotility and sperm membrane damage. We observe even earlier effects in *Prm1*-/- mice displaying ROS-mediated DNA damage already in the testis subsequently leading to immotility and disrupted membranes. Thus, loss of *Prm1* renders the ROS system more fragile at an even earlier stage. Additionally, transcriptional silencing seems impaired in *Prm1-/-* and *Prm2-/-* [33] sperm as indicated by differential gene expression analysis in testis. Further, we detected increased histone retention in *Prm1-/-* and *Prm2-/-* sperm using MassSpec. Differential abundance analysis of nuclear proteins in *Prm1-/-* and *Prm2-/-* epididymal sperm show proteins related to translation and apoptotic processes consistent with the secondary effects observed. However, only moderate differences were detected in *Prm1+/-* sperm compared to WT indicating that these changes most likely do not contribute to the phenotype observed.

While *Prm1-/-* male mice display a phenocopy of *Prm2-/-* male mice, marked differences were found between heterozygous males. Interestingly, *Prm1+/-* males are subfertile showing a reduction in average litter sizes and lower pregnancy frequencies. Of note, *Prm2+/-* are fertile [32]. This suggests that loss of one allele of *Prm1*, in contrast to loss of one allele of *Prm2*, cannot be tolerated. Transmission electron micrographs revealed that DNA of *Prm1+/-* sperm appears electron dense suggesting that the chromatin in sperm is condensed to the same level as *Prm2+/-* [32] and WT sperm. This raises the question as to why *Prm1+/-* males are subfertile.

We show that a small population of *Prm1+/-* epididymal sperm stain 8-OHdG positive. Also, genomic DNA isolated from *Prm1+/-* sperm is partially fragmented. This indicates that some sperm experience DNA damage caused by ROS rather than all sperm bearing some degree of DNA damage. Surprisingly, however, we did not detect marked differences in chromatin condensation or membrane integrity between WT and *Prm1+/-* sperm. *Prm2+/-* sperm did not show an increase in ROS-mediated DNA damage compared to WT sperm [33]. Thus, *Prm1+/-* sperm seem more sensible or more exposed to oxidative stress mediated damage compared to *Prm2+/-* sperm. This might contribute to the subfertility of *Prm1+/-* males.

Noteworthy, redox imbalance in sperm has been repeatedly connected not only to sperm DNA damage, but also reduced sperm motility in men [49]. It has been reported that sperm mitochondria present a significant source of ROS in defective sperm [50]. In human, spontaneous production of mitochondrial ROS by defective sperm causes peroxidative damage to the sperm midpiece leading to reduced sperm motility. One of the major differences between *Prm1+/-* and *Prm2+/-* sperm is that loss of one allele of *Prm1* leads to a marked decrease in sperm motility, whereas *Prm2+/-* sperm motility was not significantly different from WT sperm [32]. Only around 23% of the *Prm1+/-* sperm are motile, an amount that qualifies in human as asthenozoospermic according to the WHO criteria [49, 51, 52]. Since mitochondrial ROS has been negatively correlated to sperm motility and we detect a moderate increase in ROS in *Prm1+/-* sperm compared to WT, we believe that reduced sperm motility in *Prm1+/-* males is (at least partially) caused by ROS, which contributes to the subfertility observed in *Prm1+/-* males.

Another notable difference between *Prm1+/-* and *Prm2+/-* sperm is the aberrant DNA protamination as revealed by CMA3 staining. While approximately one third of the *Prm2+/-* sperm stain with CMA3, around 98% of *Prm1+/-* sperm are CMA3-positive. For human ejaculates, the percentage of CMA3-positive sperm varies considerably [53] and values of up to 30% CMA3-positive sperm have been defined for normal semen samples of fertile men [54, 55]. Thus, we argue that the 29% CMA3-positive sperm seen in *Prm2+/-* males, despite being higher than the values detected in WT controls, can be tolerated and do not affect regular fertility. However, it is surprising that only 2% of CMA3-negative sperm in *Prm1+/-* mice still result in a partially retained fertility. Enhanced CMA3-staining of sperm is correlated with increased histone retention. Surprisingly, the high CMA3 level in *Prm1+/-* sperm could not be correlated with increased histone retention as shown by MassSpec. One possible explanation for the intense CMA3 staining in *Prm1+/-* sperm could be the vast amounts of pre-PRM2 detected in nuclear protein extracts from epididymal sperm. We hypothesize that failure of processing of pre-PRM2 and pre-PRM2 loading onto sperm DNA might allow the intercalating dye to access DNA and stain the chromatin.

Defects in PRM2 processing were also described for histone variant H2A.L.2-KO, transition protein (TPs)-1 and -2 (TP1/TP2)-double KO, TP2-KO and cleaved PRM2 cP2-KO mouse models [40, 56-58], all of which display fertility problems. Interestingly, a recent study showed that mutation of a single non-arginine residue in PRM1 (P1^K49A/K49A^) leads to impaired PRM2 processing in mice [59]. Of note, *Prm2+/-* sperm contain scarce amounts of pre-PRM2 as well. The relative amount of pre-PRM2 is, however, significantly larger *in Prm1+/-* sperm. Hence, species-specific PRM1 levels are required for proper PRM2 processing and alterations of these levels unequivocally lead to reduced fertility. Noteworthy, presence of pre-PRM2 in subfertile human sperm has been described before [19].

In addition to high levels of pre-PRM2, *Prm1+/-* sperm display a shift in the PRM1:PRM2 ratio from approximately 1:2 in WT sperm to 1:5 in *Prm1+/-* sperm. Also, in cP2-deficient mice a shift in the protamine ratio has been described. Arévalo *et al*. have shown that mice lacking the highly conserved N-terminal part of PRM2, called cleaved PRM2 (cP2), display defective PRM2 processing and show a PRM1:PRM2 ratio of approx. 5:1 [40]. Mice lacking cP2 on one allele are infertile. As *Prm1+/-* males are subfertile, it appears that a ratio of 1:5 can be tolerated to some extent, whereas a 5:1 ratio is incompatible with fertility. Of note, the protamine ratio in *Prm2+/-* sperm is not significantly different from WT sperm explaining their regular fertility. In human, alterations of the species-specific ratio both on protein and transcript level have been repeatedly correlated to male sub- and infertility[15-27]. These results once again underline the importance of the protamine ratio in species expressing both protamines.

Of note, the protamine ratio in mice harboring a C-terminally altered allele of protamine 1 (P1^K49A/K49A^) was, contrary to the *Prm1+/-* sperm analyzed here, unaltered [59]. P1^K49A/K49A^ mice are, like *Prm1+/-* mice subfertile. Interestingly, P1^K49A/K49A^ sperm show increased histone retention similar to *Prm1-/-* and *Prm2-/-* sperm which is not detected in *Prm1 +/-* sperm. Thus, the presence of one functional *Prm1* allele is sufficient for proper histone eviction.

In summary, we generated and characterized *Prm1*-deficient mice. We demonstrate that *Prm1+/-* mice are subfertile, exhibiting sperm with moderate ROS-induced DNA damage and reduced motility. Opposed to *Prm2+/-* sperm, large amounts of pre-PRM2 were detected in *Prm1+/-* sperm. While the crucial species-specific protamine ratio is maintained in *Prm2+/-* sperm, *Prm1+/-* sperm exhibit an aberrant protamine ratio. We demonstrate that *Prm1-/-* and *Prm2-/-* mice display impaired transcriptional silencing, increased histone retention and redox imbalance leading to severe sperm damage, which render males infertile. Loss of *Prm1* seemingly triggers the ROS system at an even earlier stage compared to loss of *Prm2*. By intercrossing the *Prm1*-deficient mouse line presented here to our published *Prm2*+/- model, we will next generate and analyze *Prm1+/- Prm2*+/- double heterozygous males, which will further advance our knowledge about the molecular consequences of disturbances in PRM1 and PRM2 levels and the PRM1:PRM2 ratio.

## Acknowledgements

This study was supported by grants from the Deutsche Forschungsgemeinschaft (DFG) to HS (SCHO 503/23-1), LA (AR 1221/1-1) and KS (STE 892/19-1). We thank Gaby Beine, Andrea Jäger, Angela Egert, Irina Kosterin, Anna Pehlke, Greta Zech, Barbara Fröhlich and Tania Bloch for excellent technical assistance. Protein identifications were done at the University of Bonn Core Facility Mass Spectrometry, Institute of Biochemistry and Molecular Biology, Medical Faculty, University of Bonn funded by the Deutsche Forschungsgemeinschaft (DFG) – Projektnummer 386936527. We would like to acknowledge the assistance of the University Bonn Core Facility Next Generation Sequencing (NGS) for their support.

## Author contributions

G.E.M., K.S., and H.S. conceptualized the study. S.S. generated gene-edited mice. G.E.M., J.M., L.A., S.S., A.F. and A.K. analyzed mice. G.E.M. and H.S. drafted the manuscript. All authors read and approved the final manuscript.

