## Supplementary Figure 1 for "Loss of *Prm1* leads to defective chromatin protamination, impaired PRM2 processing, reduced sperm motility and subfertility in male mice"

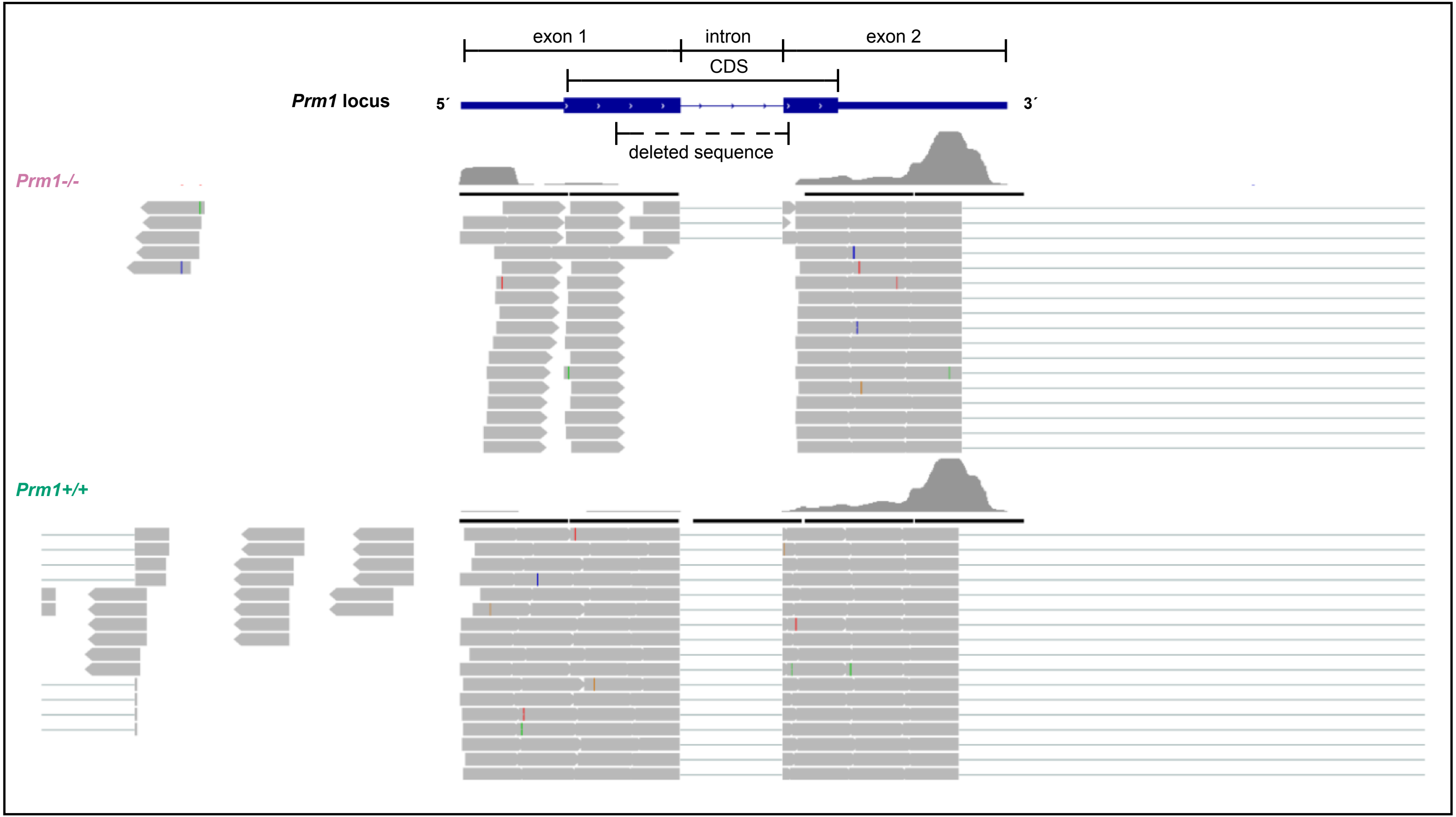


**Supplementary Fig. 1. Representative cut-out of RNAseq reads mapping to the *Prm1* locus.** Reads from whole testis RNAseq of *Prm1*+/+ and *Prm1*-/- mice.
