## Supplementary Figure 2 for "Loss of *Prm1* leads to defective chromatin protamination, impaired PRM2 processing, reduced sperm motility and subfertility in male mice"

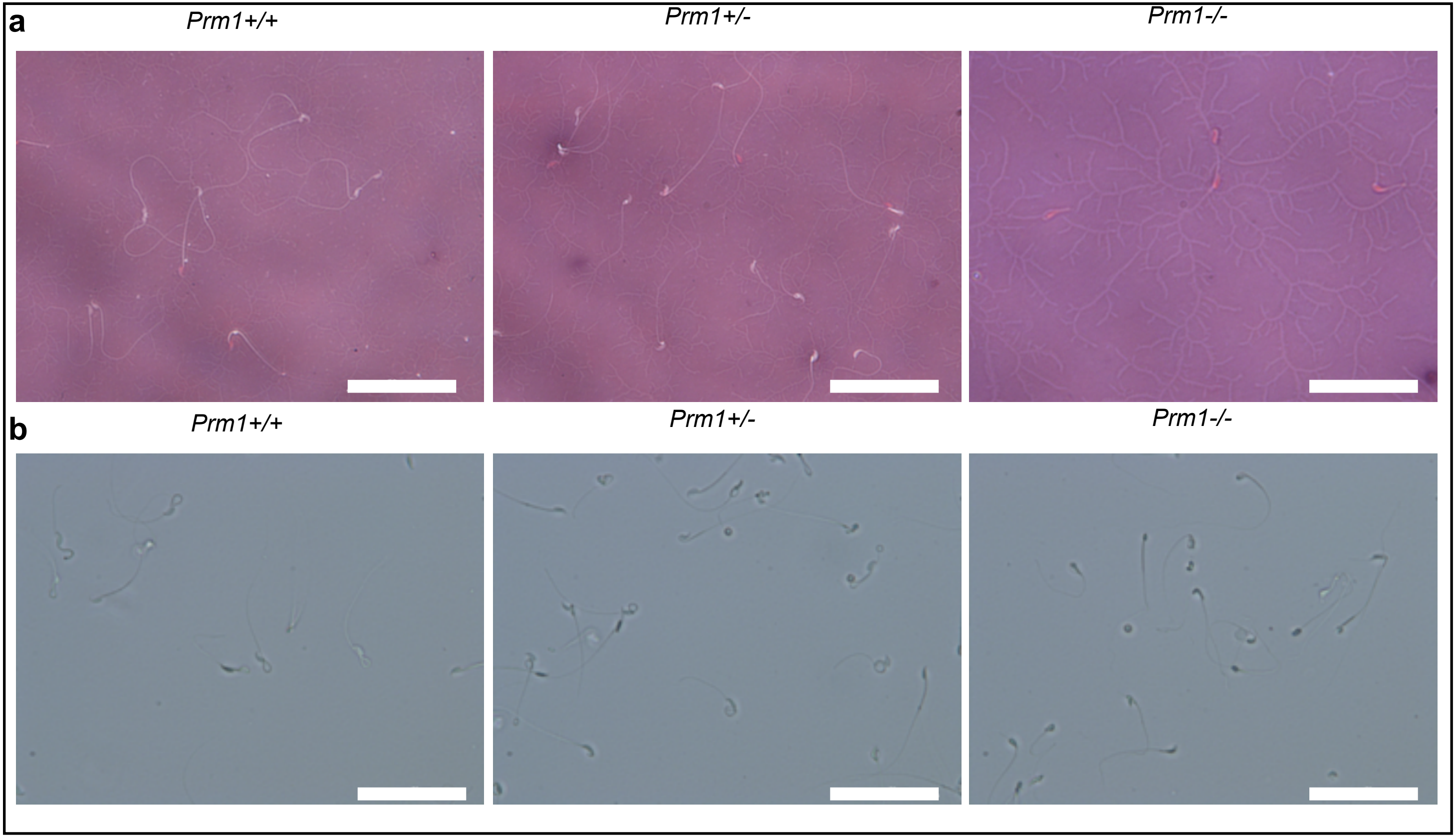


**Supplementary Fig. 2. Membrane damages of *Prm1*-deficient sperm. (a)** Representative pictures of Eosin-Nigrosin staining of *Prm1+/+*, *Prm1+/-* and *Prm1-/-* sperm. Scale: 50 µm **(b)** Representative pictures of hypoosmotic swelling tests of *Prm1+/+*, *Prm1+/-* and *Prm1-/-* sperm. Scale: 50 µm
