## Supplementary Figure 3 for "Loss of *Prm1* leads to defective chromatin protamination, impaired PRM2 processing, reduced sperm motility and subfertility in male mice"

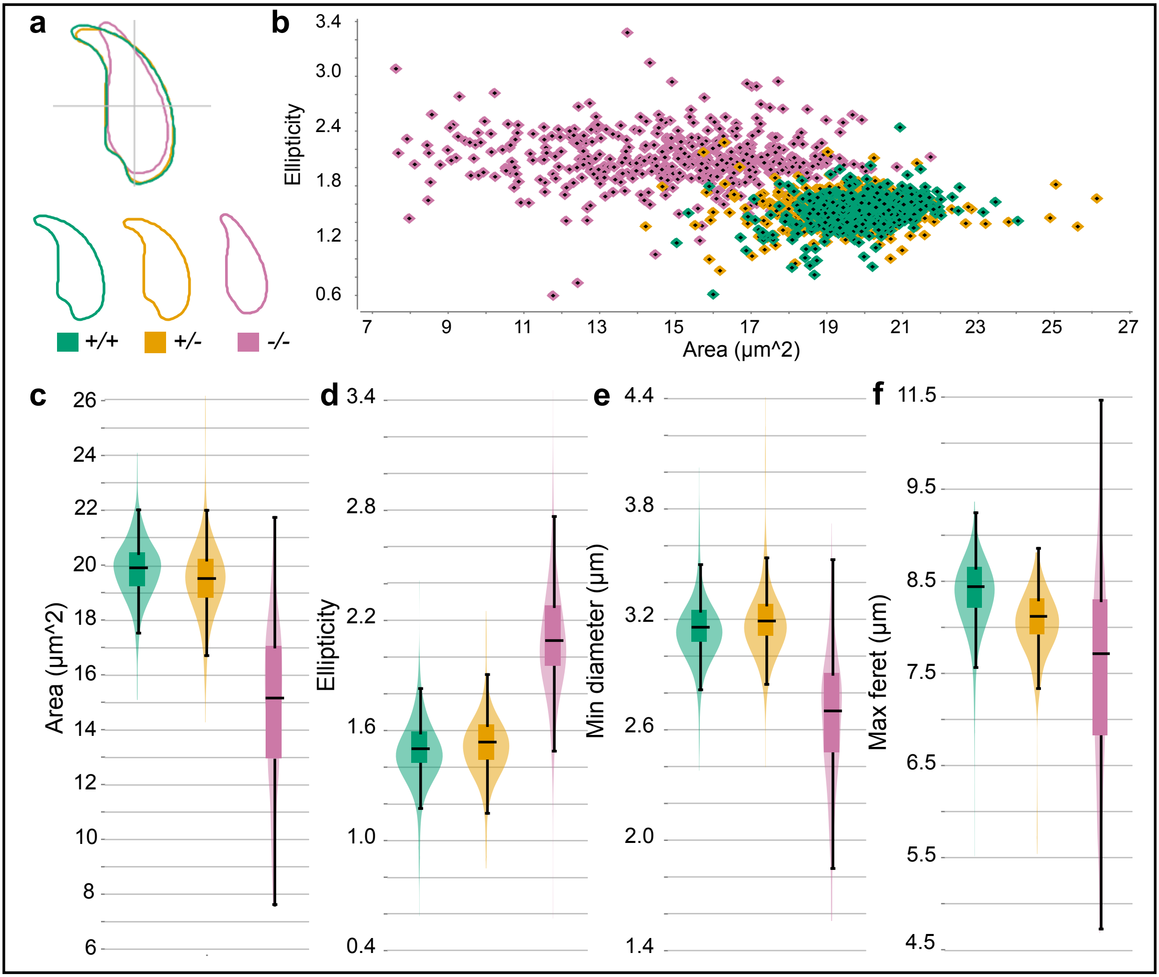


**Supplementary Fig. 3. Nuclear head morphology analysis of epididymal *Prm1*-deficient sperm. (a)** Consensus head shapes for epididymal sperm from *Prm1+/+*, *Prm1+/-* and *Prm1-/-* animals are depicted. **(b)** Scatter plot depicting the sperm head shapes of *Prm1+/+*, *Prm1+/-* and *Prm1-/-* mice by area (µm^2) and ellipticity (bonding height/ bonding width). **(c)** Violin plot presenting the mean area (µm^2) of *Prm1+/+*, *Prm1+/-* and *Prm1-/-* sperm heads. **(d)** Violin plot showing the mean ellipticity (bonding height/ bonding width) of *Prm1+/+*, *Prm1+/-* and *Prm1-/-* sperm heads. **(e)** Violin plot showing the mean minimum diameter (µm) of *Prm1+/+*, *Prm1+/-* and *Prm1-/-* sperm heads. **(f)** Violin plot depicting the mean maximum ferret (µm) of *Prm1+/+*, *Prm1+/-* and *Prm1-/-* sperm heads.
