## Supplementary Figure 4 for "Loss of *Prm1* leads to defective chromatin protamination, impaired PRM2 processing, reduced sperm motility and subfertility in male mice"

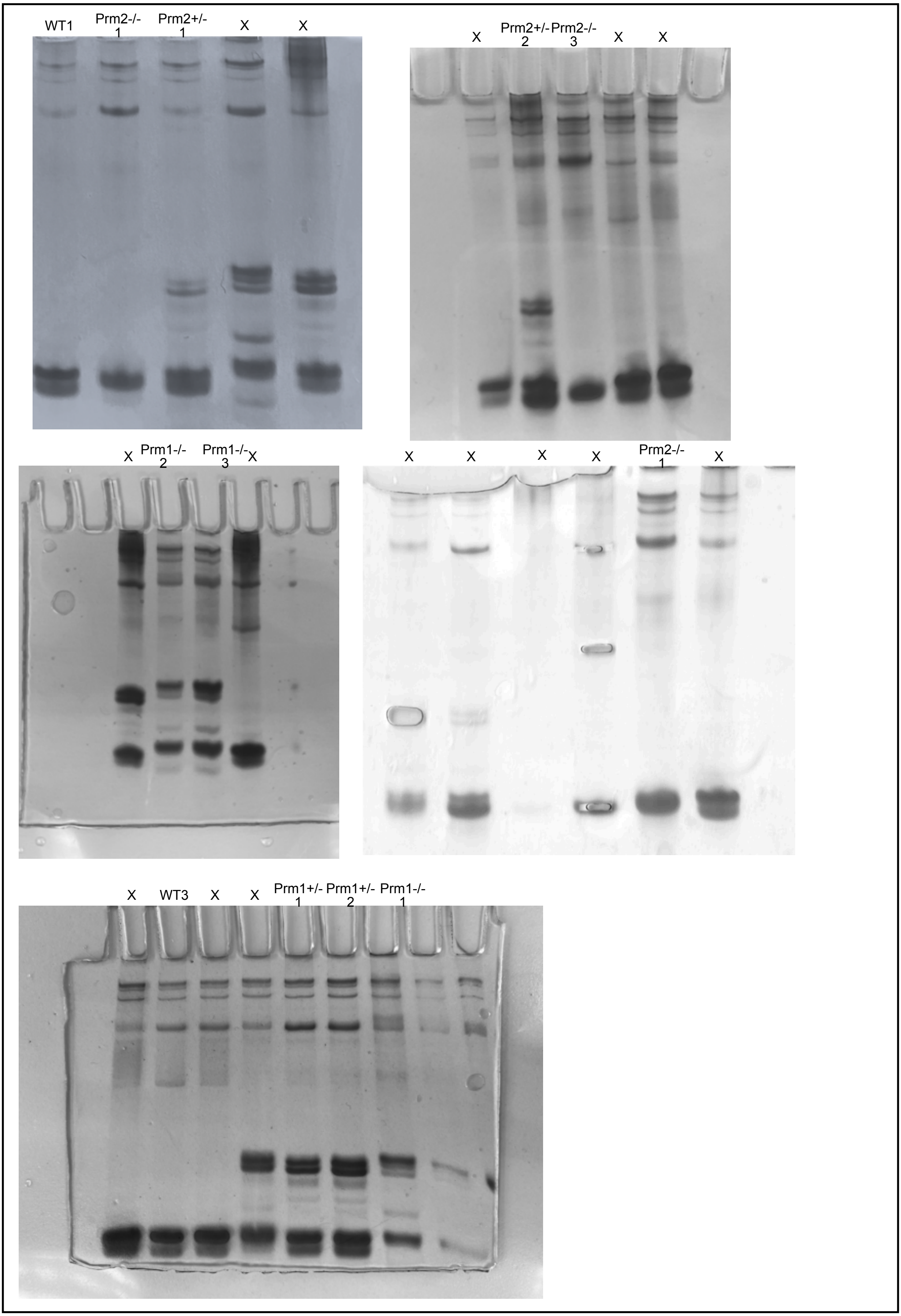


**Supplementary Fig. 4. Acid-urea gels used for band area quantification.** Lanes marked with X where not used or part of other studies.
